# Neurochemically-evoked activity in slice preparations of the octopus arm nerve cord

**DOI:** 10.1101/2025.11.29.691307

**Authors:** Gabrielle C. Winters-Bostwick, Caleb J. Bostwick, Robyn J. Crook

## Abstract

Octopus arms contain circuits that support local sensorimotor integration and autonomy, responsible for fast and flexible behaviors, but their population dynamics remain unknown. The axial nerve cord (ANC) is a series of sucker-associated ganglia whose cortex layer houses multiple classes of intermingled neurons. Here we use calcium imaging in *ex vivo* slices from arms of *Octopus bocki*, to visualize how these networks respond to controlled application of neurotransmitters and neuromodulators. Glutamate and dopamine are dominant excitatory drivers that activate most neurons in the ANC cortex, with a substantial overlapping populations and smaller transmitter-specific subsets. Glutamate continues to excite additional neurons at higher concentration, whereas dopamine responses saturate. Serotonin alone evokes mixed responses but, when applied first, consistently reduces glutamate– and dopamine-driven activation, revealing a state-dependent modulatory role. GABA and octopamine yield weak heterologous effects, and high-dose acetylcholine sharply suppresses global ANC neuronal activity while inducing muscular contraction, consistent with inhibitory cholinergic receptors in arm ganglia. Across conditions, responsive neurons show no evidence of spatial structure, with no clear anatomical segregation by transmitter response profiles. These results provide the first link between neurochemical architecture and real-time firing dynamics in a semi-autonomous circuit.

## Introduction

Octopuses exhibit behavioral flexibility and neuronal complexity that is unparalleled among invertebrates. Their large, multi-lobed brains and elaborate peripheral nervous systems underlie remarkably dynamic camouflage and mimicry, refined predatory strategies and sophisticated problem-solving^1–7^, capturing the fascination of a broad spectrum of scientists. In particular, their set of eight semi-autonomous arms is a lineage-specific innovation requiring both neural integration with their centralized brains, and decentralized coordination within and across each arm, independent of hierarchical control from the brain.

The axial nerve cord (ANC) of each arm is a series of bead-like ganglia connected to one another via aboral cerebrobrachial tracts (CBT) of axons that transmit and integrate neuronal signals locally, along the length of each arm, and up to the subesophageal brain^8–10^. Of the approximately 500 million neurons in the octopus nervous systems, 60-70% (300–350 million) reside within their arms. Each ganglion and its surrounding circuitry are associated with a single sucker and bear highly conserved structural motifs from the arm base to the tip^5^.

Cytoarchitecture in general is also largely conserved along the ANC with each ganglion composed of a dense cortex layer (CL) of somata projecting inward to a neuropil densely packed with axons and synapses. Each axial ganglion is capable of integrating external stimuli for near-instantaneous motor control, while simultaneously contributing to actions executed by the whole animal. This highly distributed organization makes octopus arms unique models for understanding how peripheral circuits can produce complex behavior.

Recent studies^11^ have revealed that the CL contains primarily small (8-12 µm diameter), densely packed cholinergic and glutamatergic neurons, with loosely stratified monoaminergic (dopaminergic, serotonergic, and octopaminergic) and peptidergic (FLRIamide and bradykinin-like neuropeptide) populations of neurons (soma diameter 8-20< µm), sparse GABAergic cells, and putative glia. Many of the cells show complex patterns of co-expression and cotransmission. Monoaminergic cell types show a pronounced base-to-tip molecular gradient, with distinct expression profiles in proximal versus newly added distal ganglia. Longitudinally, ANC neurons are organized into repeating segments separated by septa where ANC nerves exit, establishing a modular neuropil architecture and a spatial topographic map for each sucker^5,12^. The extraordinary complexity of this repeated structure, paired with constant arm growth and the addition of new ganglia at the growing tip of the arm, suggests that the octopus may have evolved unique ways to effect local network control that must remain consistent across distance from the central brain, as well as along developmental and functional gradients.

Despite decades of interest in the octopus arm’s capacity for fine motor control and distributed processing of information, we still lack a mechanistic paradigm for how ANC circuits function as ensembles to produce the complex, dynamic and exquisitely coordinated behaviors produced by the suckers, arms and body of octopuses. Recent advances in calcium imaging methods for cephalopods^13–15^ now make it possible to examine circuit properties of local arm networks as they respond to controlled applications of stimuli. Although genetically-encoded calcium indicators (GECIs) are not available in cephalopods, synthetic, high-sensitivity indicators such as Cal-520 AM have proven effective for two photon recordings in octopus optic lobes^15^. These recent studies reveal robust maps of stimulus evoked activity, shedding completely new light on the convergent principles of visual processing in evolutionary distant species. Here we adapt a similar method to the arm to capture how ANC networks respond to mechanical and chemical cues that octopuses naturally encounter.

Our functional hypotheses stem from studies suggesting broad conservation of patterns of neural activation and transmission in molluscs, but also numerous cephalopod-specific specializations. Glutamatergic and cholinergic cells are densely packed and intermingled in the cortical layer of the ANC^11^. Both neurotransmitters are established as drivers of fast synaptic transmission in cephalopod nervous systems, yet both demonstrate functional duality depending on the circuit context. Glutamate is widely recognized as a principal excitatory transmitter in cephalopod CNS, where glutamatergic synapses and receptors underlying long-term potentiation (LTP) in the vertical lobe memory circuit support and mediate excitatory visual processing in the optic lobe^16–21^. Conversely glutamate also mediates inhibition at photoreceptor terminals and contributes to complex, mixed excitatory-inhibitory signaling in the optic lobes^20^. Similarly, acetylcholine serves as the primary excitatory transmitter at the neuromuscular junction, essential for arm motor control^10,22,23^, yet it exerts inhibitory effects in sensory systems such as the statocyst^24^ and optic lobes^14^. Given the high density of these two neuronal populations in the arm, and their established capacity for both excitation and inhibition, we predicted that exogenous application of glutamate and acetylcholine would induce rapid, robust responses in the arm ganglia, though we remained agnostic regarding whether the net population activity would be excitatory or inhibitory. The role of gamma-aminobutyric acid (GABA) is less straightforward. Across invertebrates GABA often mediates rapid inhibition^25,26^, but can be depolarizing or excitatory in circuits where chloride gradients differ from canonical adult states^27^. GABA’s functional role in the ANC circuitry remains ambiguous, and its extreme scarcity in cell bodies within the ANC^11^ suggests that its influence may be exerted by a small, specialized local population and/or delivered through descending input from the central brain where larger populations of GABAergic neurons have been observed^28,29^. Finally, we hypothesized that monoamines, found in loosely organized strata of large scattered neurons in the ANC^11^, act primarily as neuromodulators that shape circuit state and plasticity^30^ but may also function as primary transmitters^14^. Monoamines like serotonin, dopamine, and octopamine have been implicated in modulation of learning and memory^28,31^, camouflage activity^32^, feeding circuitry^33–36^, and visual processing^37–39^ in central circuitry of cephalopods and in other molluscs. Recent work^14^ in the octopus visual system further shows that dopamine can exert a direct excitatory influence in central circuits, suggesting that monoamines in cephalopods can combine classic modulatory roles with more immediate control over neuronal activity.

Previous efforts to record activity in the ANC have focused mostly on identifying spiking activity in response to mechanical or chemical cues, or correlating firing to muscle activation^40–42^. Here, for the first time we use calcium imaging in slice preparations of whole arms to examine ensemble responses to perfusion with putative neurotransmitters and neuromodulatory substances. Using thick slices with intact muscle, suckers and skin, we imaged responses of hundreds of cells per slice while preserving at least some local circuitry, essential in a system where extensive processing is likely distributed through dense, unorganized and poorly characterized neuropil.

Overall, this work links neurochemical architecture to real time function in complex peripheral circuitry by combining Cal-520 AM population imaging with complementary model-based and epoch-based analyses that quantify responses in acute slices of octopus arm. We aim to define how transmitter-diverse ANC circuits encode neurochemically relevant cues, and what these dynamics imply about distributed circuit control in the highly complex and understudied peripheral nervous system of octopuses.

## Results

Our experiments provide the first real time visualization of neural population activity in octopus arms. Using calcium imaging in arm cross sections from *Octopus bocki* (Fig. 1a,b), we monitored activity across the axial nerve cord ganglion and adjacent local circuits, including intact musculature, suckers, and epithelia, within each slice. Across 57 twenty-minute recordings, these ANC preparations showed robust spontaneous firing (Fig. 1c-e) as well as diverse stimulus evoked responses to sequential chemical cues (Fig 1f,g). We summarized these population level dynamics using GLM based classifications, mean population profiles, and excitation/inhibition activity profiles, and then used these measures to compare how different neurotransmitter activate and modulate ANC networks.

**Figure 1.**
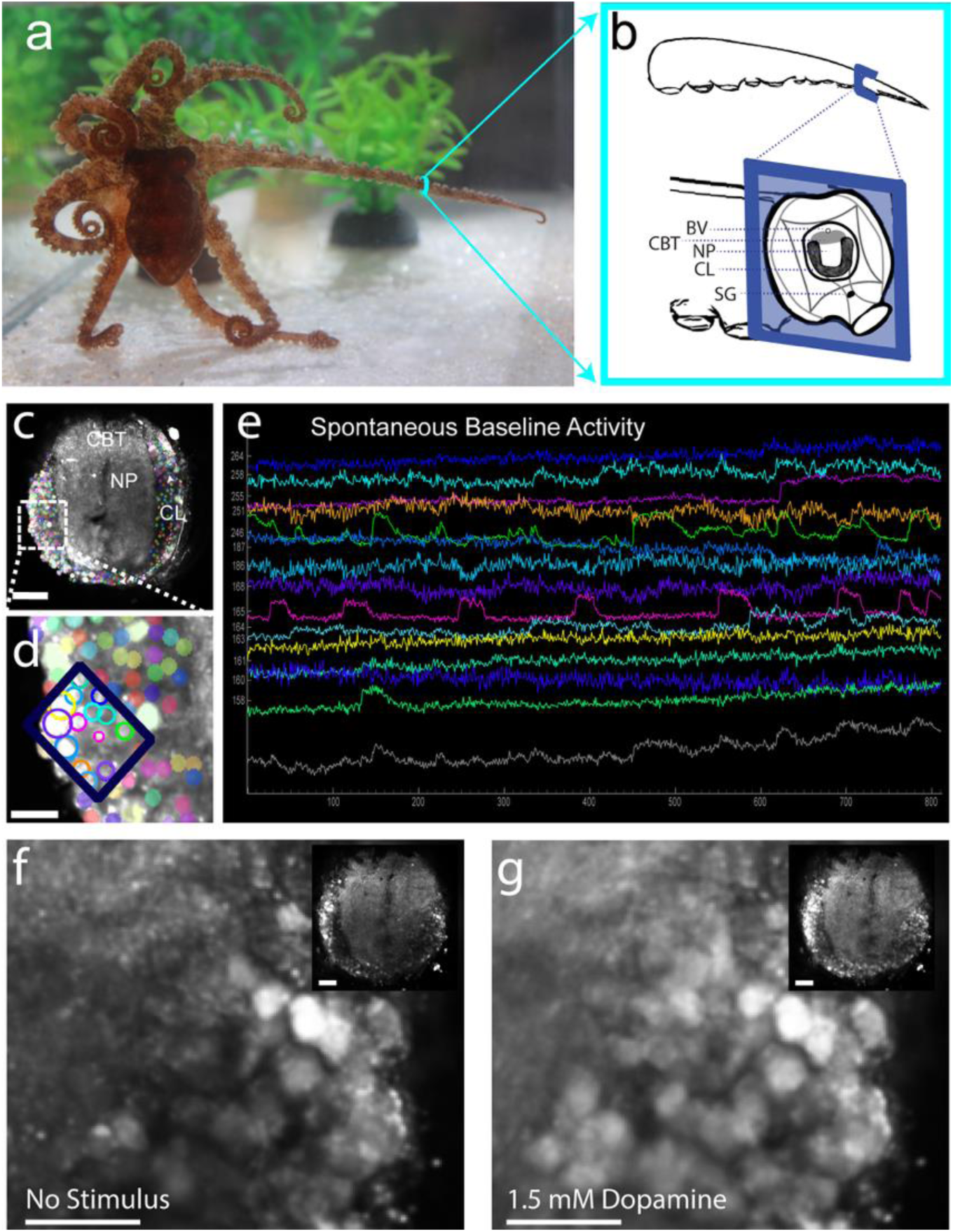
*Octopus bocki* ANC axial nerve cord preparation and example neuronal recordings. (a) Adult *Octopus bocki* used for *ex vivo* arm axial nerve cord (ANC) imaging experiments. (b) Schematic of the arm and ANC cross-section depicting the ANC ganglion region and imaging plane. (c) Suite2p segmentation output for a representative imaging field of view of the ANC showing neuronal ROIs. (d) ROI map highlighting a selected subset of neurons used for trace display in panel e. (e) Example ΔF/F traces from the selected cells in panel d during control ASW delivery, illustrating spontaneous activity patterns. (f) Mean-projection image of neurons in the ANC cortex layer (CL) after ASW negative control. (g) Mean-projection image after addition of 1.5 mM dopamine in the same preparation, illustrating marked change in fluorescence intensity in a subset of neurons in the CL. BV: Blood Vessel; CBT: Cerebro-brachial tract; NP: Neuropil; CL: ANC cortex Layer, SG: Sucker Ganglion. Scale bars: 50 µm for whole-slice panels (c, f, g) and 20 µm for further magnified panels (d).

Glutamate (Glut) and dopamine (DA) were the dominant excitatory drivers in these preparations (Figs. 2–3). When applied at a 500 µM concentration, Glut excited a large fraction of ANC neurons (Fig. 2r-Glut low→Glut high: 55.4% “E” (E=excited); Fig. 2z-Glut→DA: 75.2% E; Fig. 3r-Glut→ Serotonin (5HT): 62.8% E), all above ASW excitation baseline fractions. Dopamine at 500 µM similarly produced widespread excitation when presented first (Fig. 2j-DA low→DA high: 57.7% E; Fig. 2ah-DA→Glut: 65.4% E; Fig. 3j-DA→5HT: 67.4% E), exciting roughly two-thirds of neurons, again far above baseline (Fig. 2h). In both cases, per-cell heatmaps and MPP traces showed rapid population activation after stimulus onset (MPP traces, fig 2x and 2af for glutamate-first paradigms; Fig 2p and 2an for dopamine-first paradigms; Fig. 3r for glutamate-first with serotonin; Fig 3p for dopamine-first with serotonin), and response maps localized excited neurons distributed across the recorded population. Each transmitter also activated a notable unique subset in the two-drug datasets: in Glut→DA, a large shared pool remained excited by both stimuli (Fig. 2af-EE 50.2%), alongside a substantial Glut-only excitatory group (E0 23.3%) and a smaller Dopamine-only group (0E 7.8%); in the reciprocal DA→Glut sequence, the EE fraction was similar. (Fig. 2an-EE 50.1%) with Dopamine-only (E0 13.7%) and Glut-only (0E 15.6%). These patterns indicate transmitter-specific excitatory pools in addition to a large shared population.

**Figure 2.**
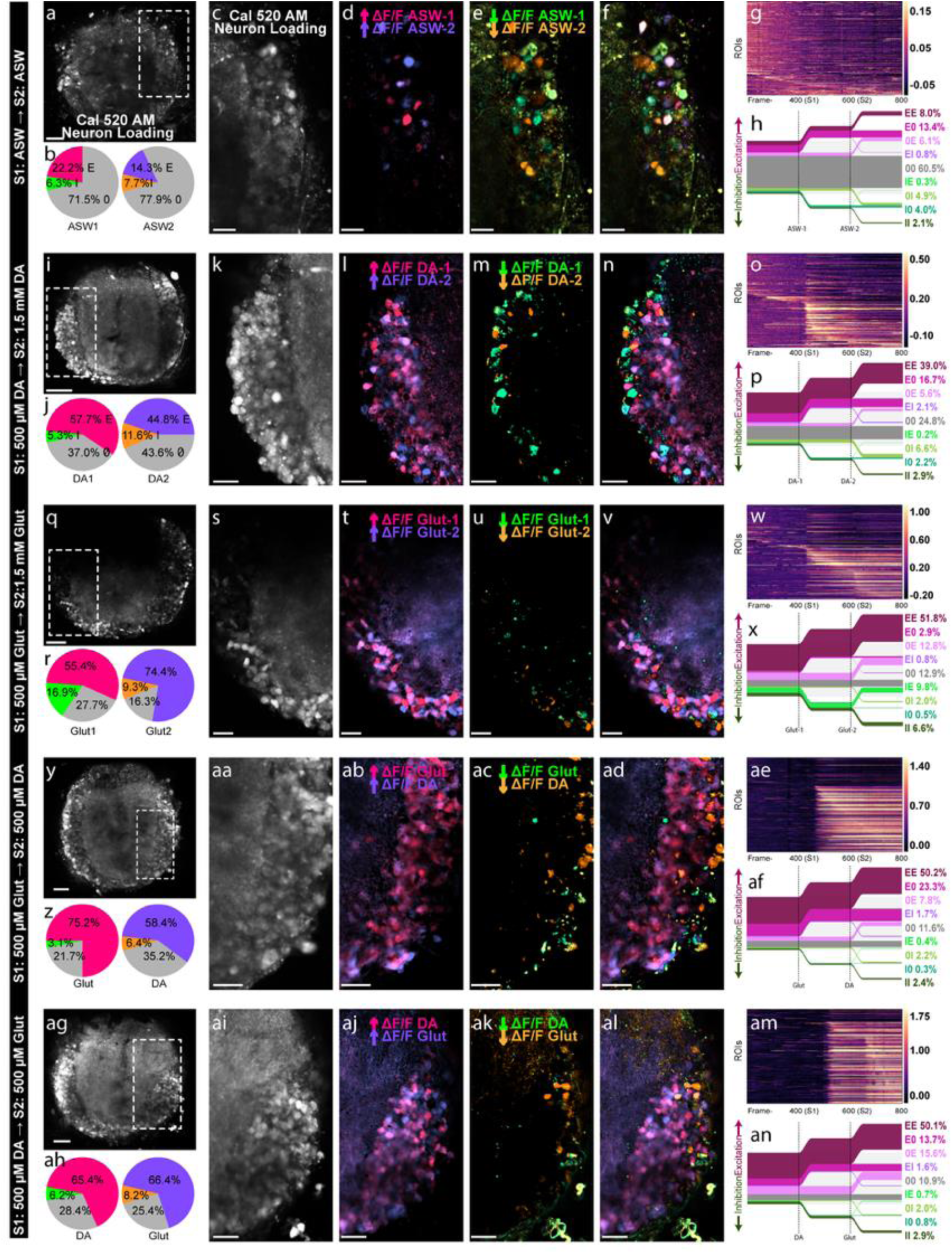
Activation and inhibition of ANC neurons by Glutamate and Dopamine. Each row of panels depicts a representative experiment from each experimental condition. Lefthand bar indicates the stimulus order for each panel row. (a, i, q, y, ag) Cal-520 AM loading for the entire imaged ganglion preparation. (b, j, r, z, ah) Response pie charts for each chemical stimulus (S1 or S2), summarizing the fraction of ROIs classified as activated (fuchsia and purple), inhibited (green and orange), or non-responsive (gray) for S1 (left) and S2 (right) pie. (c, k, s, aa, ai) Magnified mean-projection images focused on the CL neurons. (d, l, t, ab, aj) Excitation maps of increased ΔF/F in response to S1(fuchsia) and S2 (purple). (e, m, u, ac, ak) Inhibition maps of decreased ΔF/F in response to S1 (green) and S2 (orange). (f, n, v, ad, al) Overlaid maps of previous two panels. (g, h; o, p; w, x; ae, af; am, an) Per-cell heatmaps (g, o, w, ae, am) and mean population profile (MPP) traces (h, p, x, af, an) aligned to each stimulus onset illustrating stimulus-locked activation of ANC neurons by each chemical stimulus. MPP trace abbreviations: “E”-excitation/ ↑ΔF/F; “I”-inhibition/↓ΔF/F; “0” no change in ΔF/F. Scale bars: 50 µm for whole-slice images (a, i, q, y, ag) and 20 µm for remaining magnified insets.

**Figure 3.**
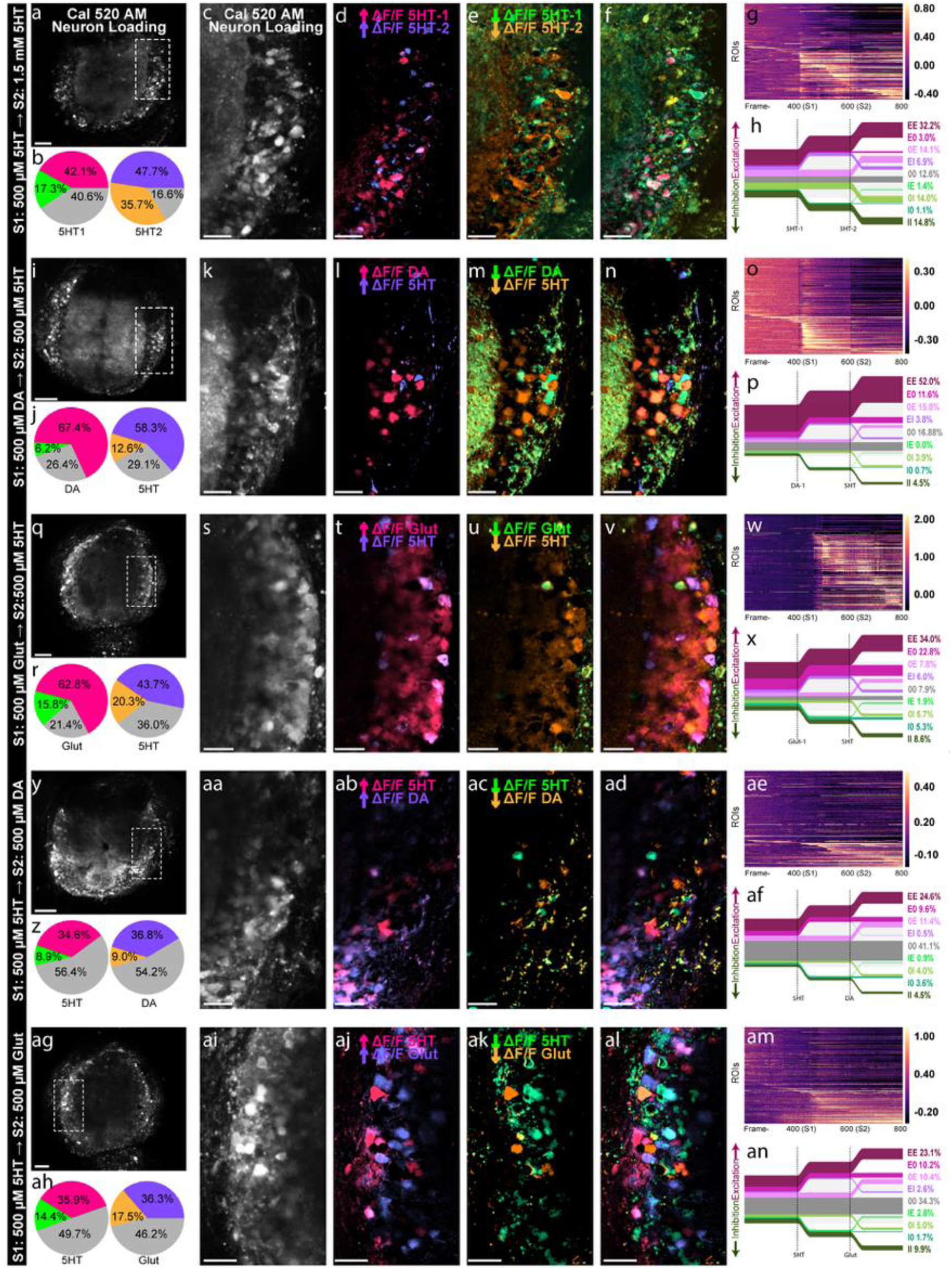
Serotonin (5HT) induces excitatory and inhibitory neural responses in ANC neurons and modulates subsequent excitatory transmitter-induced activation. In all panel rows 5HT is delivered alone or before or after an excitatory transmitter (DA or Glut). Panels are organized as in Fig. 2. Lefthand bar indicates the stimulus order for each panel row. (a, i, q, y, ag) Cal-520 AM loading for the entire imaged ganglion preparation. (b, j, r, z, ah) Response pie charts for each chemical stimulus (S1 or S2), summarizing the fraction of ROIs classified as activated (fuchsia and purple), inhibited (green and orange), or non-responsive (gray) for S1 (left) and S2 (right) pie. (c, k, s, aa, ai) Magnified mean-projection images focused on the CL neurons. (d, l, t, ab, aj) Excitation maps of increased ΔF/F in response to S1(fuchsia) and S2 (purple). (e, m, u, ac, ak) Inhibition maps of decreased ΔF/F in response to S1 (green) and S2 (orange). (f, n, v, ad, al) Overlaid maps of previous two panels. (g, h; o, p; w, x; ae, af; am, an) Per-cell heatmaps (g, o, w, ae, am) and MPP traces (h, p, x, af, an) aligned to each stimulus onset illustrate that S1 delivery of 5HT reduces S2 activation by excitatory neurotransmitters (DA and Glut). Scale bars: 50 µm for whole-slice images (a, i, q, y, ag) and 20 µm for remaining magnified insets.

Dose-response comparisons revealed distinct excitation properties for glutamate versus dopamine (Fig. 2). In paired glutamate-only experiments, the second glutamate application at 1.5 mM consistently increased excitation relative to the initial 500 µM dose (Fig. 2r and Fig. 2x-S1: 55.4% E; S2: 74.4% E), with a clear S2-only excitatory cohort and a rise in total S2 excitation (Fig. 2x-0E 12.8%). This pattern suggests heterogeneity in glutamate activation thresholds across ANC neurons, with additional cells engaged only at higher concentration or after repeated exposure. In contrast, dopamine dose-series recordings showed no comparable expansion of the excitatory pool at 1.5 mM (Fig. 2j and 2p-S1: 57.7% E; S2: 44.8% E). Instead, excitation plateaued at the lower dose and often declined on the second application, with limited S2-only activation (Fig. 2p-0E 5.6%). Together, these results indicate saturating dopamine activation by 500 µM, whereas glutamate continues to excite additional neurons at higher dose.

Sequential dopamine and glutamate experiments showed that these transmitters activate both shared and unique neuronal subsets, without strong order-dependent interaction (Fig. 2). In Glut→DA recordings, initial application of glutamate was robustly excitatory, and subsequent dopamine application still activated a large excitatory population (Fig. 2z and 2af-S1: 75.2% E; S2: 58.4% E). The reciprocal DA→Glut sequence produced a similar profile structure (Fig. 2ah and 2an-S1: 65.4% E; S2: 66.4% E). Overall, both sequences elicited strong excitation by each transmitter with comparable excitation patterns. The persistence of both shared and unique neuronal cohorts across orders, and the stable EE fraction, suggests that glutamate and dopamine act in parallel excitatory pathways within ANC circuitry rather than in a strictly hierarchical arrangement.

Serotonin responses were variable in sign and magnitude, but serotonin strongly altered subsequent network excitability (Fig. 3). When serotonin was delivered as S2 after glutamate or dopamine, many neurons still exhibited excitation above ASW S2 baseline (Fig. 3r and 3x-Glut→5HT S2: 43.7% E; Fig. 3z and Fig. 3af-DA→5HT S2: 58.3% E), and EE profiles remained common (Fig. 3x-EE 34.0%; Fig. 3af-EE 52.0%), indicating a sizeable shared excitatory pool with these transmitters.

However, when serotonin was delivered first, subsequent excitation by glutamate or dopamine was consistently suppressed (Fig. 3ah and 3an-5HT →Glut S2: 36.3% E; Fig. 3z and Fig. 3af-5HT →DA S2: 36.8% E), despite remaining above baseline. Comparisons of the proportion of cells excited when glutamate was delivered first vs. when glutamate was applied after serotonin showed that this difference was significant (unpaired t-test, t=3.31, p=0.0095). The effect was slightly weaker although still significant for application of dopamine, which also showed reduced excitation of cells when applied after serotonin, compared to application without a prior stimulus (unpaired t-test, t=2.92, p=0.02). This suppression was accompanied by a clear increase in inhibition above ASW S2 baseline (Fig. 3p S2: 22.9% “I”, inhibition) and elevated non-responsive profiles (Fig. 3an and 3p-“00” fraction 34.3% and 41.1%) with reduced EE activation (Fig. 3an and 3p-EE 23.1% and 24.6%). Thus, serotonin appears to function as a potent modulator of circuit state, lowering the likelihood that ANC neurons will be subsequently activated by excitatory transmitters.

Other transmitters produced weaker or predominantly inhibitory effects that further underscore transmitter-specific tuning of ANC networks (Fig. 4). GABA induced inhibition that exceeded ASW baselines and scaled with dose (Fig. 4b and Fig. 4h-S1: 16.8% I; S2: 28.8% I), consistent with graded inhibitory control rather than broad suppression. Octopamine elicited comparatively small and inconsistent deviations from baseline, with many neurons remaining non-responsive across doses (Fig. 4j and Fig. 4p). A high acetylcholine concentration (1.5mM) showed the most pronounced inhibitory shift, where high-dose ACh drove widespread inhibition above baseline with very limited excitation in the Cal-520 AM channel (Fig. 4r,x-S2: 86.6% I and 7.3% E), indicating strong transmitter-specific engagement of inhibitory pathways. Concurrent red-channel recordings (AlexaFluor 568 loading dye; Fig. S1o–ab) show no visible pre/post intensity loss (Fig. S1o–r) and no evidence of global optical artifacts (Fig. S1s–v), confirming that the near-global suppression in the green channel reflects true activity changes rather than bleaching or relaxation of tissue out of the imaging focal plane. Although the red-channel pie charts (Fig. S1z) include small fractions labeled as excitation or inhibition, the red-channel heatmap displays an order-of-magnitude lower ΔF/F variance with no temporal structure (Fig. S1aa–ab), indicating that these classifications reflect low-amplitude noise around a flat baseline.

**Figure 4.**
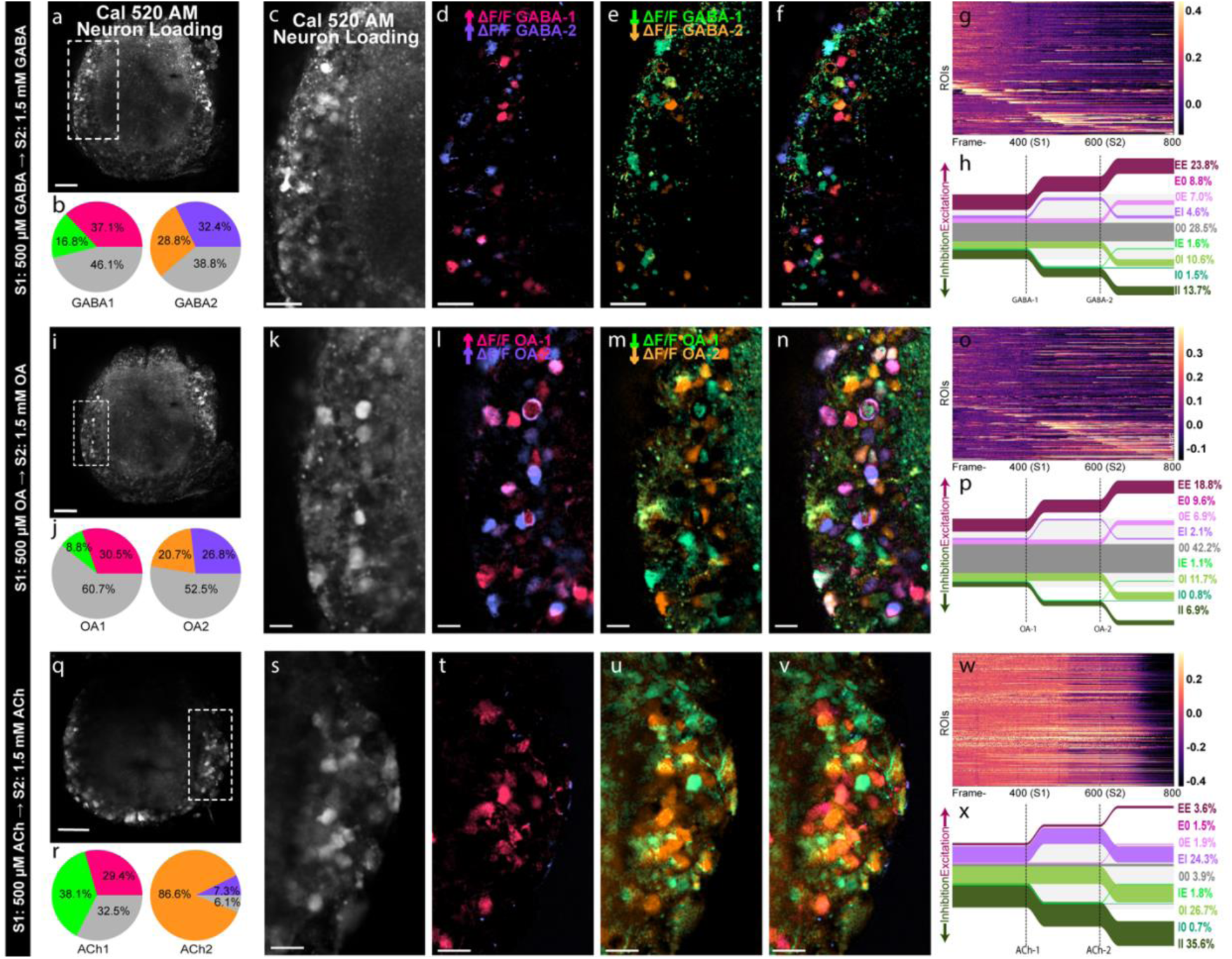
GABA and octopamine induce variable neural responses and acetylcholine suppresses globally. Panel organization matches Fig. 2. Lefthand bar indicates the stimulus order for each panel row. (a, i, q) Cal-520 AM loading for the entire imaged ganglion preparation. (b, j, r) Response pie charts for each chemical stimulus (S1 or S2), summarizing the fraction of ROIs classified as activated (fuchsia and purple), inhibited (green and orange), or non-responsive (gray) for S1 (left) and S2 (right) pie. (c, k, s) Magnified mean-projection images focused on the CL neurons. (d, l, t) Excitation maps of increased ΔF/F in response to S1(fuchsia) and S2 (purple). (e, m) Inhibition maps of decreased ΔF/F in response to S1 (green) and S2 (orange). (f, n, v) Overlaid maps of previous two panels. (g, h; o, p; w,x) Per-cell heatmaps (g, o, w) and MPP traces (h, p, x) aligned to each stimulus onset illustrating stimulus-locked activation of ANC neurons by each chemical stimulus. Scale bars: 50 µm for whole-slice images (a, i, q) and 20 µm for remaining magnified insets.

Anatomical response maps in Fig. 5a (for Glut, DA, 5HT, OA, GABA, and ACh) show that responsive ROIs are spatially intermingled across the layer of neuronal somata rather than forming discrete anatomical strata, indicating no detectable topographic organization of transmitter-activated neuronal cohorts in the cross-sectional plane. Together, Fig. 5a demonstrates that transmitter effects observed in GLM contrasts reflect broad, distributed excitement patterns with distinct temporal signatures, rather than localized populations of chemically tuned neurons.

**Figure 5.**
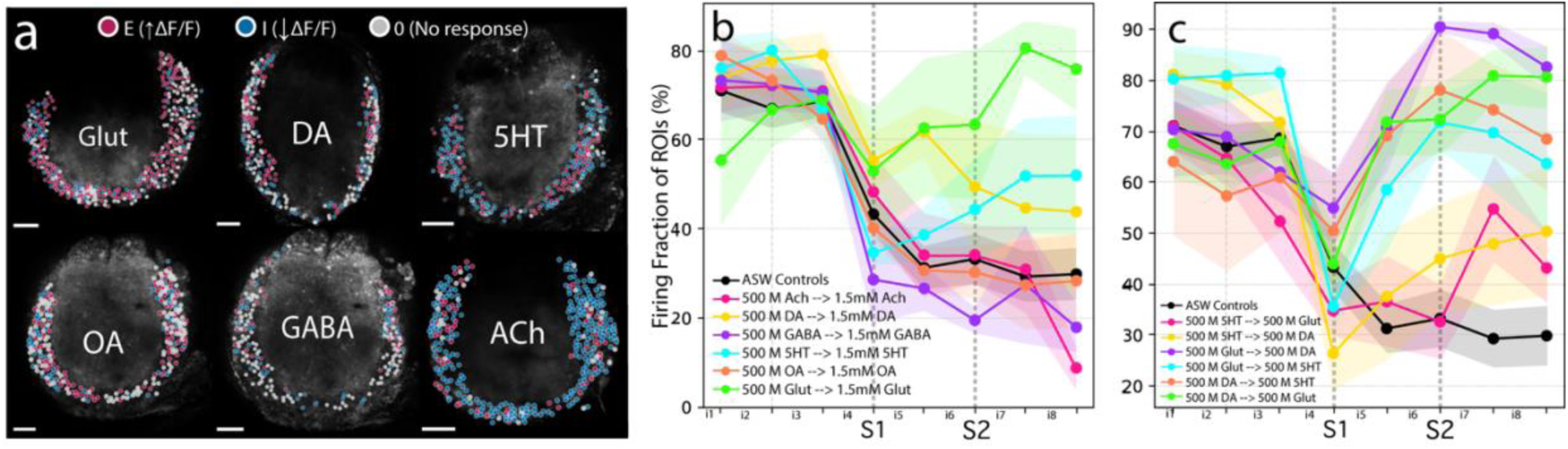
Neurochemical identity and sequence induce distinct ANC neuron activation profiles without clear spatial pattern. (a) Representative anatomical maps of responsive ROIs for each transmitter condition (Glut, DA, 5HT, OA, GABA, ACh), illustrating the absence of any detectable anatomical organization of neuronal ensembles based on chemical stimulus responses. (b) Dose-dependent effects of neurotransmitter exposure. Overlay plot of mean ± SEM fire fraction over time in dose-increase experiments; each line represents a stimulus condition, with ASW controls in black. (c) Equivalent fire-fraction overlay for same-dose transmitter-switch experiments, again including ASW controls in black. Fire fraction here is defined as the percent of cells that fired (crossed a per-cell noise-based threshold) at least once within each interval, which differs from the GLM analyses in earlier figures that directly contrast stimulus windows against baselines. Stimuli were applied after acclimatization and stabilization of baseline firing (visualized as a dropoff in fire fraction after interval 3). Scale bars: 50 µm.

In the dose-increase experiments (Fig. 5b) all six neurochemicals tested show distinct temporal and dose-dependent activity profiles. Glut drives strong stimulus-locked activation above ASW controls, with activity rising after S1 and increasing further after high-dose S2, consistent with the S2-only excitatory cohort seen in the MPPs (Fig 2x). DA likewise shows robust excitation after the 500 µM application but no further increase following high-dose S2, matching the saturating dopamine dose response inferred from GLM contrasts and MPP classifications (Fig. 2p). 5-HT produces modest and variable activation at 500 µM, but shows a clearer upward shift at the higher dose, aligning with heterogeneous serotonin effects that strengthen with concentration. Notably, because fire fraction counts only positive events, any mixed excitation and inhibition within a paradigm will be collapsed into a single “active” fraction, whereas the GLM MPPs resolve these bidirectional cohorts. The effects of OA exposure remain weak across intervals, with activity staying near control levels and showing no clear dose-dependent excitation, consistent with minimal OA effects in the MPPs (Fig. 4p). GABA induces relatively low activity during low dose stimulus intervals and a subtle overall downward shift at higher dose. Finally, a high dose of ACh produces the strongest suppressive signature: activity collapses in the final interval, paralleling the near-global inhibition detected in the Cal-520 channel (Fig. 4q-x) and reinforcing that ACh induces a dominant inhibitory shift at high doses. Together, Fig. 5b reinforces transmitter-specific dose properties identified in GLM analyses.

Transmitter-combination experiments (Fig. 5c) extended these findings to sequential interactions among excitatory and modulatory cues. Both combinations of DA and Glut show strong increases in activity after each stimulus and sustained engagement across intervals, consistent with a large shared excitatory pool plus transmitter-specific subsets and no detected order-dependent interference. In contrast, serotonin-first paradigms (5HT→Glut and 5HT→DA) show reduced excitation by the second (excitatory) transmitter relative to the corresponding Glut-first or DA-first sequences, independently confirming that serotonin reduces subsequent excitatory activity. Overall, Fig. 5c complements the GLM/MPP analysis by demonstrating that excitation by glutamate and dopamine is stable, while serotonin acts as a strong modulator on later population engagement.

Fire fraction analyses (Fig 5b-c) provide an independent, time-resolved corroboration of the GLM/MPP results by tracking the fraction of neurons that cross a per-cell activity threshold across eight equal intervals spanning the full 20-min recording. Because fire fraction is based on suprathreshold positive ΔF/F events, it tracks population engagement over time (as opposed to in comparison to a baseline) and can reveal suppression as a reduction in event occurrence, but it does not explicitly quantify inhibitory (negative) shifts captured by the GLM/MPP analysis.

## Discussion

This study links neurochemical architecture to circuit dynamics in the decentralized nervous system of octopus arms by characterizing real-time dynamics of chemical stimulus-induced activity and modulation in highly heterogenous neuronal populations in octopus arm ANCs. Molecular and anatomical work have mapped the locations of diverse cell types and structures in the ANC^11^, characterized the ultrastructure of repeating segmented ganglia^5^, and generated sucker-oriented topographic maps along the arm axis^12^, but we had yet to visualize population activity at rest or during physiologically-relevant chemical input. Using calcium imaging in slice preparations of the whole arm, we observed how neurons of the ANC cortex respond as a network, while preserving aspects of their anatomical setting. Our approach builds on recent calcium imaging studies in the optic lobe^14,15^ by adapting similar methods to peripheral neural tissue that retains muscle, chromatophores, suckers, and sensory epithelia. We observed that glutamate and dopamine reliably served as strong excitatory transmitters, activating large, overlapping sets of neurons in the ANC cortex with little evidence for hierarchical gating between them. Serotonin behaved primarily as a state-setting modulator that amplified glutamatergic and dopaminergic excitation rather than driving large responses on its own. High-dose acetylcholine produced robust suppression of ongoing activity, and GABA and octopamine exerted only weak or inconsistent effects under our experimental conditions. Together, these patterns indicate that octopus arm ganglia support flexible chemical control through parallel excitatory and modulatory pathways operating within broadly intermingled neuronal networks, providing a first functional bridge between ANC neurochemistry and circuit-level dynamics.

### Dopamine and glutamate act as robust excitatory neurochemicals in ANC ganglia

Both dopamine (DA) and glutamate (Glut) evoked strong and consistent activation of CL neurons relative to ASW control. Although monoaminergic neurons are far less abundant than the ubiquitous intermingled glutamatergic and cholinergic populations in the ANC CL^11^, our results suggest that DA, alongside glutamate, exerts a major excitatory influence on peripheral arm neurons. DA’s strong excitatory signature here suggests a direct role in shaping arm-ganglion output as opposed to a purely modulatory function. This interpretation is consistent with recent studies in the optic lobe, where DA can act as an excitatory transmitter and neuromodulator in visual circuits^14^, and with work in gastropod sensorimotor systems showing that DA can exert both fast excitatory and inhibitory actions in addition to slower modulatory roles.^43^.

Our results reveal a pronounced overlap between DA– and glutamate-responsive populations, potentially by acting on a shared postsynaptic pool or by inducing release of Glut by the numerous glutamatergic neurons. The latter option suggests presynaptic modulation, a common monoaminergic mechanism in molluscs^28,44,45^. At the same time, the presence of ANC neurons reported to co-secrete DA and glutamate^11,39,46^ offers an additional angle: a single presynaptic population could deliver fast glutamatergic excitation while co-released DA tunes response magnitude, duration, or threshold in a state-dependent way, naturally yielding high overlap without requiring two independent inputs. We also noted that increasing glutamate concentration continued to excite additional CL neurons at higher doses, whereas DA-evoked activation tended to plateau, suggesting that glutamatergic drive can progressively engage larger portions of the local network while dopaminergic excitation saturates.

These data also raise the possibility of very short-loop glutamatergic recurrence, including potential autapses. Because glutamatergic neurons are abundant in the CL, bath-applied glutamate could be exciting the numerous CL neurons that themselves release glutamate, including via recurrent local connections or direct self-synapses, amplifying activation through a positive-feedback loop. We cannot resolve this with our current recordings, but it would explain why glutamate produces such robust population-level activation and why DA, if co-released or acting presynaptically on glutamatergic terminals, tracks that same cohort so closely. The smaller DA-only and glutamate-only responder sets are therefore most parsimoniously interpreted as reflecting differences in receptor expression, synaptic architecture, position within local circuitry, rather than clean segregation into separate, non-overlapping circuits.

### Serotonin modulates excitatory neurotransmitter activity

Serotonin (5HT) alone produced highly variable population effects across recordings, which may reflect differences across septate ganglion segments, slice boundaries, or baseline state. Segment-to-segment heterogeneity is a plausible contributor because ganglia are modular, and different segments can be captured in different slices even within the same ganglion^5,12^. The strong monoaminergic gradients described from arm base to tip^11^ further suggest that these arm tip slices from newly added distal ganglia may inherently show weaker or more labile serotonergic signatures than those from the arm base. Despite this variability, 5HT showed a consistent modulatory effect on activity of excitatory neurochemicals. When applied first, 5HT reduced subsequent DA– and Glut-evoked excitation. This pattern fits classic monoaminergic roles in molluscs and cephalopods, where serotonin often shifts circuit dynamics and behavioral state instead of acting as the primary fast excitatory molecules^32,44,45,47^. Metabotropic 5HT receptors, presynaptic gating, and disinhibitory routing are all well-established serotonin mechanisms that could explain a “network state switch” effect here^45,48^. In the context of a segmented ANC with repeating sucker-associated modules, such state-setting actions could allow monoamines to bias whole stretches of the arm toward varying degrees of responsiveness without reconfiguring local circuitry.

### GABA and octopamine effects are inconsistent at the population level

Both GABA and OA transmitters produced indistinct or modest ΔF/F changes in most datasets. One possible explanation for this is that their postsynaptic targets are sparse within the imaged CL pool. This is especially relevant for GABA, whose somata are extremely scarce in ANC molecular maps and appear restricted to medial regions of the lateral ANC CL, possibly suggesting highly localized control that could go undetected in a large cell population. Another factor to consider is response polarity, as GABA, which is mostly known for its inhibitory influence^25,26^, can be depolarizing (excitatory) in circuits where chloride gradients are non-canonical, including immature or dynamically tuned networks^27^. If chloride varies across developing tip ganglia, GABA could produce mixed excitation and inhibition that goes undetected at the population scale. Finally, regardless of GABA’s specific role in the ANC, its strongest influence may arrive via descending input from the central brain rather than local interneurons, consistent with sparse local GABAergic cells. This interpretation fits well with broader cephalopod anatomy, where more substantial GABAergic populations are located in central motor and learning centers^28,29^, potentially allowing the brain to impose “top-down” control over otherwise highly autonomous arm modules. Unlike GABAergic neurons, octopaminergic neurons are large and relatively abundant in the ANC, but the ambiguous activity of exogenous OA application does not point compellingly toward any functional interpretation. It’s possible that OA is restricted to a neuromodulatory role, and that our experimental paradigms did not recapitulate an environment where OA would be exerting a large enough modulatory effect at a detectable spatial or temporal scale.

### Acetylcholine diminishes neuronal activity at high doses

Acetylcholine produced a dose-dependent suppression of ANC activity that we interpret as genuine inhibition. Lower concentrations yielded weak or mixed effects, whereas high-dose ACh reliably induced a large drop in CL somatic calcium signals. Concurrent red-channel imaging implies this is not a bleaching artifact or shift in focal plane caused by large changes in muscle tone, but a true reduction in excitability or calcium influx compared to baseline spontaneous activity. This pattern fits a growing body of work showing that ACh can act as an inhibitory transmitter via the anion-selective AChRB receptors in coleoid^14^ and other lophotrochozoan^49,50^ circuits. In the octopus optic lobe, ACh gates the AChRB1 receptor, which is expressed in photoreceptors and dopaminergic amacrine cells, and acute ACh application predominantly suppresses activity in these networks^14^. Together with earlier physiological work showing ACh-mediated inhibition at octopus photoreceptor terminals^51^, these findings support the idea that multiple cholinergic synapses in coleoid circuits are inhibitory rather than excitatory.

In our preparation, bath-applied ACh reliably triggered large, rapid contractions of residual arm musculature surrounding the slice at both experimental doses, indicating that the neuromuscular junction remained strongly responsive and that ACh is almost certainly excitatory in octopus arm muscles^10,22,23^. Conversely, somatic calcium activity in CL neurons was strongly suppressed, consistent with ACh simultaneously activating inhibitory AChRB-like receptors on CL neurons. Functionally, the combination of excitatory action at the neuromuscular junction and suppression within the ganglion compartment may suggest a mechanism by which a strong cholinergic input can contract the arm while an inhibitory cholinergic “brake” in the ANC prevents prolonged contraction states. Regardless of the precise mechanisms, we observed that elevating ambient ACh can strongly silence peripheral ANC network activity, in line with a broader role for cholinergic signaling as a circuit-level control of cephalopod sensory and motor pathways^14,51^.

### Alternative Organizational Principles in the Arm ANC

The absence of any detectable anatomical organization of neurochemical-induced responses in the ANC CL suggests intermingled transmitter coding in shared CL populations. Rather than mapping onto strictly segregated circuits, responsive pools overlapped across all transmitters, sometimes even in the same cell as in DA and Glut. That implies that ANC modules may use shared neuronal substrates whose activity is tuned by transmitter-specific dynamics, supporting flexible local integration necessary for arm autonomy. In the context of ANC architecture, multiplexed coding in shared CL ensembles would allow the same structure to participate in different behaviors like grasping, exploration, or withdrawal, depending on the present combination of neurotransmitters in each neuron’s microenvironment. These functional findings are consistent with our molecular study, which found only loose strata of enriched monoaminergic and peptidergic cell types and no laminar or clustered organization of glutamatergic and cholinergic neurons. In many vertebrate and invertebrate systems, lamination, modular clustering, lateral inhibition, and distinct ON and OFF pathways act as near-universal strategies to reduce the complexity of coding and stabilize circuit function^52–55^. Recent work on the octopus optic lobe suggests that this structure follows many of those same information-processing rules, including clear lamination, contrast-enhancing antagonistic surrounds, and separate pathways for light-on and light-off signals ^15,39,56^. In comparison, the lack of obvious structural rules in either the molecular or functional architecture of the arm ANC is striking, and suggests that arm circuits may rely on fundamentally different organizational principles to achieve flexible, autonomous control.

### Methodological Considerations and Future Directions

Several methodological constraints in this exploratory study limit how far we can interpret some of the collected data. Low frame rate and droplet-based delivery without controlled perfusion mean we cannot resolve spike-like kinetics, precise stimulus onset, or true dose–response relationships; our “low” versus “high” conditions should be treated as coarse comparisons. Likewise, the patterns recorded at these low frame rates represents summed activation across multiple cells within each circuit, and we are not able to discriminate between immediate postsynaptic activation vs. larger network initiation as a result of our stimulus applications. Variability in dye loading, variable animal age and unknowable health status, and slice position along the arm likely adds noise, especially for transmitters with subtle or state-dependent actions such as 5HT, GABA, and octopamine. In any slice preparation, some network connections are inevitably severed and likewise our slices do not fully recapitulate the ANC’s natural mechanical and neuromodulatory environment.

A promising next step would be to dissect these pathways at the receptor level with more precise stimulus control and timing. Antagonists and receptor-specific tools could distinguish direct from polysynaptic effects, separate fast ionotropic actions from slower metabotropic ones, and identify which presynaptic versus postsynaptic targets best explain the order– and state-dependence we observe. Higher-framerate, perfused, segment-resolved imaging should reveal local, mixed-sign modulation that is invisible in our current population view. In parallel, comparing proximal and distal ganglia, and imaging across known segment boundaries, will be essential for understanding how developmental gradients and modular arm organization shape transmitter activation along the ANC. Complementary studies at the connectomic and transcriptomic level could also help resolve fine details of the organization of the axial nerve cord.

Despite these limitations, our data clearly demonstrate that ANC circuits in distal arm ganglia are strongly excitable through glutamate and dopamine, while serotonin reliably tunes excitatory gain in subsequent excitatory neurotransmitter applications, and acetylcholine can drive population-level suppression at high concentrations. This functional picture complements prior work showing that computation in octopus arms is local, modular, and tightly linked to specific behaviors^10,12,41,42,57–59^, further underscoring the ANC as a powerful site for connecting transmitter architecture to arm autonomy. Here, we have begun to link transmitter architecture to the real-time physiology of a peripheral circuit that supports octopus arm autonomy and integration, and it sets up a tractable framework for dissecting how neuromodulatory context shapes decentralized control in cephalopod nervous systems.

## Methods

### Octopus Acquisition and Housing

Adult *Octopus bocki* were acquired from Sea Dwelling Creatures (Los Angeles, CA) between February and June, 2025. Octopuses were housed individually in three-gallon enclosures within a larger 1600 L recirculating artificial seawater system at 23.5-25.5 °C. Recirculating artificial seawater for housing (hASW) was prepared by dissolving Instant Ocean™ Sea Salt in reverse osmosis purified water per manufacturer’s instructions and filtered through physical, chemical, and biological systems. Octopus enclosures were enriched with shells, rocks, plastic plants, PVC pipes, and each octopus was fed 1-2 live grass shrimp per day. Animal health was monitored daily.

Ethical Note: Octopuses are invertebrates and therefore are excluded from regulatory oversight in the USA; thus, no IACUC protocol was required for this study. However, we adhered to Directive 2010/63/EU^60^ and ARRIVE guidelines for characterizing standards of care, humane endpoints, and experimental procedures.

### Octopus Euthanasia and Dissection

Ten adult *O. bocki* were anesthetized, euthanized, dissected, and injected in “Anesthetic MgCl₂ Solution” (AMS): experimental artificial sea water (ASW) (460 mM NaCl, 10 mM KCl, 10 mM glucose, 10 mM HEPES, 55 mM MgCl₂, 11 mM CaCl₂, pH 7.4-7.6) mixed with an equal volume of 330 mM MgCl₂ in RODI water, at 13–15 °C. All solutions (ASW and AMS) were kept aerated using an air stone and portable pump through the duration of the experiment.

Once the octopus submerged in AMS became unresponsive to a mantle pinch, the animal was transferred to a dissection dish in AMS and promptly decerebrated before any further processing. The arm crown was removed and half of the arms were immediately transferred to 50 mL of ASW and stored at 4°C for experiments on the following day (repeated using the same protocol as below). The remaining arms were prepared immediately for injection by cutting ∼0.5-1.5 mm slices in the transverse or cross-sectional plane along the most distal 1/3 of the arm (toward the tip). All tissues surrounding the axial nerve cord including skin and muscle were included in the slices, and at least one sucker was left intact per slice.

### Dye Loading

Cal-520 AM (AAT Bioquest Cat. #21130) (1.3 μg/μL) and Alexa Fluor 568 Hydrazide (Thermo Fisher Scientific Cat. #A10437) (0.25 μg/μL) were dissolved into ASW containing 10% Pluronic F-127 ^∗^20% solution in DMSO (AAT Bioquest Cat. #20052) and pressure injected directly into the neuropil of each slice using a glass micropipette needle (Harvard Apparatus Cat# 30-0038) attached to a pressure injection pump (μPUMP, World Precision Instruments).

After injection, slices were embedded in 4% low-melt agarose (Sigma-Aldrich Cat. #A0701-25) and kept in aerated ASW for 30-60 minutes before imaging to allow complete cell loading of calcium dye. Only ANC slice preparations that displayed robust spontaneous baseline neural activity after the initial acclimation period were advanced to chemical stimulus recordings. See Table S1 for key resources.

### Chemical stimlui preparation

Reagents for chemical stimuli were all prepared fresh daily as 1 M stock solutions in ASW. Serotonin hydrochloride (5-HT; Thermo Scientific Chemicals, Cat# B21263.03), dopamine hydrochloride (DA; Thermo Scientific Chemicals, Cat# A11136), L-glutamic acid monosodium salt (glutamate; Thermo Scientific Chemicals, Cat# J63424.14), acetylcholine chloride (ACh; Thermo Scientific Chemicals, Cat# L02168.14), γ-aminobutyric acid (GABA; Thermo Scientific Chemicals, Cat# A11016.22), and (±)-octopamine hydrochloride (octopamine; Thermo Scientific Chemicals, Cat# J61281.03) were obtained from Thermo Scientific. See Table S1 for key resources.

### Calcium imaging recordings and stimulus protocol

All twenty-minute recordings (n=57; 48 slices from 10 octopuses) in this study adhered to a consistent structured format. The initial acclimation period (5 minutes) allowed neurons to reach a baseline of spontaneous activity after tissue relocation. After acclimation, we applied an initial bath of ASW (no chemical stimulus) to negate unintentional mechanical stimulation upon chemical application. The recorded activity during the five minutes between application of the ASW negative control and the first chemical stimulus was used as the baseline for downstream fire fraction analysis and single chemical dose experiments. At minute 10 the first chemical stimulus (Stimulus 1, S1) was applied, and at minute 15 a second stimulus (Stimulus 2, S2) was applied. Recordings were terminated at minute 20. For ASW-only negative control recordings, ASW droplets were applied at minutes 5, 10, and 15 using the same timing, and these runs were processed with the same GLM and fire-fraction pipelines and treated as single-drug recordings in which ASW occupied both stimulus positions.

Fluorescent images were acquired using a Leica Stellaris DMI8 Confocal microscope with immersion Obj. HC APO L 10x/0.30 W U-V-I (Leica Cat # 11506142) and Las X 4.6.0.27096 software or a Zeiss LSM 710 Confocal microscope with ZEN Microscopy Software and W Plan-Apochromat 20x/1.0 DIC M27 75mm objective (Zeiss Cat# 421452-9800-000). On the Leica Stellaris, Diode 405 was used to acquire DAPI images and WLL (85% of max power) was used to acquire all other channels. All images were taken in a 1024×1024 pixel format at 0.68Hz for Leica Stellaris images, and 0.64Hz for Zeiss LSM 710 images. Images were exported in TIFF format for analysis.

## DATA ANALYSIS

**Z-correction** was performed on each recording using a 0.95 frame-to-median image correlation-coefficient cutoff in the custom Matlab script (zbinCorr.m) generated for calcium imaging analysis in optic lobe by the laboratory of Dr. Cris Niell^15^.

**Custom Python script analyses were used for all subsequent processing. Script parameters are in Tables S1-S2**.

### Suite2p motion correction, segmentation, and trace extraction

Preprocessing was run in Suite2p v0.14.4 on single-plane, single-channel TIFF movies with both rigid and non-rigid motion correction, followed by combined functional (sparse) and Cellpose-assisted anatomical ROI detection, neuropil extraction, and built-in deconvolution; all key parameters and their values are listed in the Suite2p Settings Summary table. Two Suite2p acquisition/extraction settings were overridden downstream in Python: Suite2p’s assumed sampling rate (fs=10 Hz) was replaced with an effective 0.68 Hz for all temporal conversions, and Suite2p’s neuropil coefficient (neucoeff=0.7) was adjusted (effective α=0.5).

### Calcium imaging data import and ROI selection

For each session, Suite2p^61^ outputs were loaded in Python^62^, including somatic fluorescence traces (F.npy), neuropil traces (Fneu.npy), ROI inclusion flags (iscell.npy), ROI centroids/statistics (stat.npy), and acquisition metadata (ops.npy). Analyses were restricted to manually curated ROIs labeled as cells (iscell[:,0] == 1).

### Preprocessing and ΔF/F calculation

All time-to-frame conversions used a fixed, manually specified sampling rate (0.68Hz for Leica Stellaris images, and 0.64Hz for Zeiss LSM 710 images). Neuropil subtraction was enabled, yielding an effective neuropil coefficient α = 0.5. Slow baseline drift correction and global signal regression were selectively enabled based on the dye loading efficiency and the observed amount of photobleaching per recording.

Baseline fluorescence (F0) was computed per ROI as the 20th percentile of corrected fluorescence within the interval from Stimulus 1 onset to Stimulus 2 onset. ΔF/F was calculated as (F − F0)/F0.

### Stimulus timing and labeling

The ASW droplet at minute 5 and subsequent S1 and S2 chemical applications at minutes 10 and 15 were logged for each recording and used as the canonical stimulus onset times for all downstream analyses. Stimulus onset frames were extracted from a per-run stimulus summary CSV by matching the run identifier to the post-z-correction stimulus frame number. Stimulus labels were parsed from the run ID using a stimulus-code mapping (e.g., ASW and subsequent chemical cues), and repeated stimuli were suffixed to preserve ordering.

### GLM-based response detection using FIR kernels

This model reports whether cells show reliable, time-locked changes in ΔF/F during defined stimulus windows, relative to baseline. Stimulus responsiveness was quantified with a finite-impulse response (FIR) general linear model using explicit baseline-contrast designs. For single-drug dose-response experiments, we used GLM_METHOD=“explicit_baseline_contrast”, in which the baseline for all stimulus windows was defined as the frames between ASW and Stim 1. For multi-drug experiments, we used GLM_METHOD=“explicit_baseline_contrast_2drugs” to account for sustained effects of earlier chemicals, with the Stim 1 window baselined to ASW→Stim 1 frames and the Stim 2 window baselined to the frames recorded 60 seconds before Stim 2.

The FIR design matrix comprised non-overlapping rectangular bins aligned to stimulus onset with zero modeled delay. Post-onset windows were stimulus-specific: ASW responses were modeled over 60 s with 10 s bins to capture rapid mechanical-delivery effects, whereas chemical stimuli were modeled over 300 s with 30 s bins.

To control for mechanical droplet artifacts, the model applied a two-stage contrast: (1) ASW FIR bins were contrasted against a 60-s pre-ASW baseline, and (2) each chemical stimulus FIR response was contrasted against the ASW-period baseline (ASW onset to the next stimulus). Per-ROI stimulus effects were assessed with F-tests jointly across each stimulus’ FIR bins, and p-values across all ROI×stimulus tests were corrected using Benjamini–Hochberg FDR of 0.05.

### Effect-size gating and response classification

ROIs were required to pass both statistical significance and effect-size criteria. Adaptive noise-based gating was enabled with per-cell thresholds set to 3x baseline standard deviation. A fixed ΔF/F threshold of 20% was used. Among FDR-significant ROIs, responses were classified as **activated** if the largest-magnitude FIR beta coefficient was positive and exceeded threshold, **inhibited** if negative and exceeded threshold, and **unresponsive** otherwise. For each stimulus position, excitation or inhibition was interpreted as chemically evoked only when the fraction and magnitude of responsive ROIs exceeded the corresponding ASW-only control recordings.

### Mean population profiles and activity-profile classification

For visualization and summary of population dynamics, we computed mean population profiles (MPP) by averaging ΔF/F traces across all curated ROIs for each recording and stimulus condition. Based on the sign and magnitude of each ROI’s GLM-estimated response and ΔF/F trace relative to the thresholds described above, neurons were assigned to one of three activity profiles: excited (E; net increase in ΔF/F), inhibited (I; net decrease in ΔF/F), or non-responsive (0; no change). These E, I, and 0 labels were used to generate the per-stimulus pie charts and other population-level summaries shown in the figures.

### Fire-fraction analysis

In contrast to the GLM, fire-fraction summarizes within-epoch activity by asking only whether ΔF/F crosses an adaptive effect-size threshold in each epoch, without comparing responses across different epochs. Population activity was summarized as “fire-fraction,” defined as the percentage of curated ROIs that became active within each predefined interval. Fire-fraction was computed from the ΔF/F traces and curated ROI set described above, using an adaptive per-cell noise threshold. For each ROI, baseline noise (σ) was estimated from ΔF/F values **only during the ASW→Stim 1 interval**, and the activity threshold was set to 3×σ ROIs were counted as active within an interval if any ΔF/F sample in that interval exceeded their individual threshold. Fire-fraction summaries (active cell counts and percentages per interval) were used for downstream aggregation and comparison across recordings.

## QUANTIFICATION AND STATISTICAL ANALYSIS

All statistical analyses were performed in Python using SciPy (v1.13.1) and Statsmodels (v0.14.4). FIR-GLMs were fit by ordinary least squares to ΔF/F traces with stimulus-aligned regressors. Stimulus effects were assessed using F-tests across FIR bins and controlled for multiple comparisons using Benjamini–Hochberg FDR at α=0.05. Responsive ROIs were additionally required to exceed adaptive per-cell effect-size thresholds (3× baseline SD). Comparisons of proportional excitation to assess possible serotonergic modulation were conducted in Graphpad Prism 10.0. Unless otherwise noted, summary values reported in the main figures are means across recordings.

### Image projection and difference-map analyses

Directional difference images and overlays were computed from saved baseline, Stim1, and Stim2 projections (MAX or MEAN selectable via ––source-type) using *maxproj.py*. Projections (created using FIJI^56^ by importing motion corrected tiffs and exporting MAX or MEAN projections from selected windows) were loaded as float32 grayscale arrays. Four positive-only directional differences were computed by clipping negatives to zero:

### Stim1−Baseline (EXC1), Stim2−Stim1 (EXC2), Baseline−Stim1 (INH1), and Stim1−Stim2 (INH2)

Optional class-specific thresholds were applied prior to visualization using fractional cutoffs relative to the mean intensity of each projection pair. Excitatory difference maps were left unthresholded, whereas inhibitory maps were filtered to retain only changes ≥25% of the paired-image mean in order to account for unintentional photobleaching. All directional differences were normalized for display via percentile clipping (1st–99.9th) and colorized with a unified palette, then merged into EXC-only, INH-only, and ALL overlays using additive layer-blending.

Signed bidirectional maps were additionally generated from unclipped differences using symmetric robust scaling to the 99.5th percentile of |diff|. Bidirectional outputs were filtered to exclude low-magnitude changes below 25% of the paired-image.

## Acknowledgments

We thank Sarah Giancola-Detmering and all Crook Lab members for assistance with animal care and husbandry and insightful discussions. We thank SFSU’s Cell and Molecular Imaging Center (CMIC) and Dr. Annette Chan for extensive assistance with confocal microscopy and access to equipment, specifically the Leica Stellaris 5 funded by NSF MRI 2018239. We also thank the laboratory of Dr. Cris Niell, including Dr. Judit Pungor, Angelique Allen, and Dr. Anne Liu for extensive assistance with calcium imaging techniques and analysis. This study was funded by an Allen Distinguished Investigator Award to R.J.C. in Neural Circuit Design, from the Frontiers Group of the Allen Foundation and NSF IOS-2047331, to R.J.C.

## Author contributions

Study design, G.C.W.-B. and R.J.C.; experiments, G.C.W.-B.; data collection, G.C.W.-B. data analysis, G.C.W.-B. and C.J.B.; figure preparation, G.C.W.-B. and R.J.C.; writing – original draft, G.C.W.-B., C.J.B., and R.J.C.; writing – review & editing, G.C.W.-B., C.J.B., and R.J.C.; project management, G.C.W.-B. and R.J.C.; funding acquisition, R.J.C.

