## Supplementary material for "Neurochemically-evoked activity in slice preparations of the octopus arm nerve cord": Fig. S1

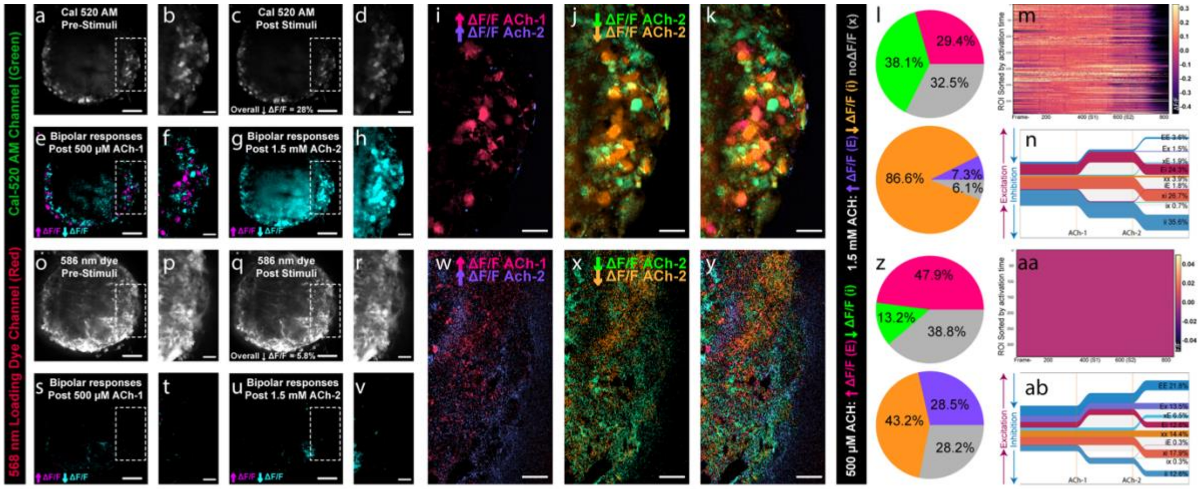

**Figure S1. High-dose Acetylcholine (ACH) suppresses activity in axial nerve cord neurons**

All data are from a single selected ACh dose-increase experiment. Concurrent recordings in the green (Cal-520 AM; a–n) and red (AlexaFluor 568; o–ab) channels contrast true cellular activity with possible optical artifacts that could be misattributed to neuronal responses. (a–d; o–r) Representative ANC ganglion showing neuronal fluorescence in the pre-stimulus and post-stimuli mean projection views. (e–h; s–v) Bidirectional representation of excitatory and inhibitory responses after 500 µM ACh (S1- e,f; s,t) and 1.5 mM ACh (S2- g,h; u,v)**.** In the Cal-520 AM channel, 500 µM ACh (e–f) evokes moderate decrease in ΔF/F with some instances of excitation, whereas 1.5 mM ACh (g–h) produces a near global net decrease in ΔF/F. (s-t) Bidirectional maps of the 568 nm channel (created using the same parameters as those of e–h) illustrate no evidence of global optical artifacts (no changes detected). (i–k; w–y) Excitation (i, w) and inhibition (j, x) maps of ΔF/F in response to S1(↑ΔF/F fuchsia; ↓ΔF/F green) and S2 (↑ΔF/F purple; ↓ΔF/F orange) followed by an overlay of previous panels (k, y). (l,z) Response pie charts for each chemical stimulus (S1 or S2), summarizing the fraction of ROIs classified as activated (fuchsia and purple), inhibited (green and orange), or non-responsive (gray) for S1 (top) and S2 right (bottom). (m, n; aa, ab) Per-cell heatmaps and mean traces aligned to each stimulus, summarizing the population-level shift from moderate suppression at 500 µM to global suppression at 1.5 mM in the Cal-520 channel and no detectable impacts in the 568-nm red channel (loading dye only). This channel should not respond to neuronal stimulation and shows no parallel intensity loss or spatial artifact during high-dose ACh, supporting a physiological (not precipitate/quenching) origin of the Cal-520 suppression. Scale bars: 50 µm (whole-slice) and 20 µm (magnified panels).

Table S1- Key Resources

| REAGENT or RESOURCE | SOURCE | IDENTIFIER |
|---|---|---|
| **Chemicals, peptides, and recombinant proteins** | | |
| **Cal-520 AM ester** | AAT Bioquest | Cat. #21130 |
| **Alexa Fluor 568 Hydrazide** | Thermo Fisher Scientific | Cat. #A10437 |
| **Pluronic F-127 (20% solution in DMSO)** | AAT Bioquest | Cat. #20052 |
| **Agarose, low gelling temperature Type VII-A** | Sigma-Aldrich | Cat. #A0701-25G |
| **Serotonin hydrochloride (5-HT)** | Thermo Scientific Chemicals | Cat# B21263.03 |
| **Dopamine hydrochloride (DA)** | Thermo Scientific Chemicals | Cat# A11136 |
| **L-Glutamic acid monosodium salt** | Thermo Scientific Chemicals | Cat# J63424.14 |
| **Acetylcholine chloride (ACh)** | Thermo Scientific Chemicals | Cat# L02168.14 |
| **γ-Aminobutyric acid (GABA; 4-aminobutyric acid)** | Thermo Scientific Chemicals | Cat# A11016.22 |
| **(+/–)-Octopamine hydrochloride** | Thermo Scientific Chemicals | Cat# J61281.03 |
| **Deposited data** | | |
| **Calcium imaging data** | This study | Available upon request from Lead Contact |
| **Experimental models: Organisms/strains** | | |
| ***Octopus bocki*** | Sea Dwelling Creatures (Los Angeles, CA) | N/A (no RRID assigned) |
| **Software and algorithms** | | |
| **Suite2p** | Pachitariu et al[59]., MouseLand | v0.14.4; **RRID:SCR_016434** |
| **Python** | Python Software Foundation | v3.10 |
| **NumPy** | NumPy developers | v1.26.4 |
| **SciPy** | SciPy developers | v1.13.1 |
| **pandas** | pandas developers | v2.3.1 |
| **statsmodels** | statsmodels developers | v0.14.4 |
| **scikit-learn** | scikit-learn developers | v1.6.1 |
| **tifffile** | tifffile developers | v2024.8.30 |
| **matplotlib** | Matplotlib developers | v3.9.4 |
| **Cellpose (used within Suite2p)** | Stringer et al.[61] | in Suite2p v0.14.4 |
| **MATLAB** | MathWorks | MATLAB R2021b; **RRID:SCR_001622** |
| **Custom calcium imaging analysis pipeline (*CellAnalysis.py; firefraction.py; meanproj.py; maxproj.py)*** | This study | Available upon request from Lead Contact |

Table S2. Analysis configuration and constants (Python pipeline: *CellAnalysis.py*)

| Parameter / constant | Value used | Function |
|---|---|---|
| **Input files** | Suite2p outputs (F.npy, Fneu.npy, iscell.npy, stat.npy, ops.npy) | Data and metadata per session |
| **ROI inclusion** | iscell[:,0]==1 | Restricts analysis to manually curated ROIs |
| MANUAL_FRAMERATE_HZ | **0.68 Hz or 0.64 Hz** | Frame rate for all time↔frame conversions in Python |
| ΔF/F baseline percentile | **20th percentile** | Defines per-ROI baseline F0 |
| ΔF/F baseline interval | **Stim1 → Stim2** (fallback: 0 → Stim2) | Frames used for F0 and baseline noise SD |
| USE_NEUROPIL_SUBTRACTION | **True** | Python-side neuropil subtraction |
| FORCE_NEUROPIL_ALPHA | 0.5 | Manual α override |
| NEUROPIL_ALPHA_FALLBACK | 0.5 | α if Suite2p coeff missing |
| APPLY_LOWESS_DETREND | Varied | LOWESS drift removal |
| LOWESS_FRACTION | 1 (inactive) | LOWESS span |
| USE_GLOBAL_REGRESSION | **False** | Subtracts global mean ΔF/F |
| GLM_METHOD | **"explicit_baseline_contrast" or** "explicit_baseline_contrast_2drugs", | FIR-GLM response calling mode |
| Supported GLM alternatives | "explicit_baseline_contrast_2drugs", "full_trace_intercept" | Other implemented modes |
| CHEM_LOCAL_BASELINE_SECONDS | 60 s | Local baseline duration in _2drugs mode |
| FIR delay | 0 s (all stims) | Shifts modeled response onset |
| STIM_CONFIG["ASW"] | window **60 s**, bins **10 s** | FIR basis for ASW control |
| STIM_CONFIG["default"] | window **300 s**, bins **30 s** | FIR basis for chemical stimuli |
| GLM fit | **OLS** | Estimation of β kernels |
| GLM test | **Joint F-test** across bins per stim | Detects evoked effect |
| FDR_ALPHA | **0.05** | BH-FDR significance cutoff |
| USE_NOISE_BASED_THRESHOLD | **True** | Adaptive vs fixed effect-size gate |

| | | |
|---|---|---|
| NOISE_MULTIPLIER | **3** | Gate = 3× baseline SD |
| FIXED_DFF_THRESHOLD | 0.2 | Fixed ΔF/F gate if adaptive off |
| USE_ASW_EFFECT_SIZE_CONTROL | **False** | Requires chem peak > ASW peak |
| ASW_FOLD_THRESHOLD | 1.1 | Fold-difference for ASW control |
| Response sign rule | sign of largest- | β |
| Post-stim state window | **300 s (5 min)** | Window for oscillatory/sporadic/stable state calls |
| PSD method | Welch PSD | Oscillation detection |
| Oscillation band | **0.003–0.2 Hz** | Slow rhythm band tested |
| Power ratio criterion | **≥5× median band power** | Oscillatory threshold |
| Min cycles | **≥2 cycles/window** | Prevents false oscillatory calls |
| FIRE_FRACTION_DFF_MULTIPLIER | **3.0** | Threshold for "active" in fire-fraction |
| Fire-fraction intervals | **first 100 frames**, **full trace**, **last 100 frames** | Where fire-fraction is computed |
| Exploratory metrics | noise floor; median-based SNR; population sparseness | QC / descriptive summaries |
| Embeddings (full traces) | PCA 2D; t-SNE 2D | Low-D visualization |
| Embeddings (GLM betas) | PCA 2D; t-SNE 2D; UMAP 2D | Low-D response-fingerprint space |
| Clustering distance | Hamming on categorical response vectors | Similarity of response profiles |
| Clustering linkage | Ward hierarchical | Dendrogram construction |
| Cluster cut | 0.7× max dendrogram height | Defines final clusters |
| Cluster QC | Silhouette (Hamming) | Cluster separation |
| Anatomical assignment | ROI centroids vs binary *Mask.tif | Region labels |
| Regional enrichment stats | $\chi^2$ per stim + BH-FDR | Tests region×response association |
| Optional peak classifier | RUN_PEAK_THRESHOLD_ANALYSIS=False | Non-GLM peak/trough calling |

* Suite2p parameters later overridden in Python.

Table S3. Analysis configuration and constants (Python pipeline: *firefraction.py*)

| Parameter / constant | Value used | Function |
|---|---|---|
| **Fire-fraction script** | *firefraction.py* | Population activity quantification |
| FIRE_FRACTION_DFF_MULTIPLIER | **3.0** | Fire-fraction activity gate |
| Fire-fraction threshold baseline | **ASW onset → Stim1 onset** | Frames used for σbaseline in fire-fraction |
| Fire-fraction intervals | Pre-Stim (first 100), Full, Post-Stim (last 100), First Half, Second Half, Intervals 1/8–8/8, plus **Frame1–ASW**, **ASW–Stim1**, **Stim1–Stim2**, **Stim2 onward** | Interval set for fire-fraction summaries |
| Projection script | *meanproj.py* | Mean-projection construction |
| Difference/overlay script | *maxproj.py* | Directional diffs, overlays, bidirectional maps |
